# Cold-induced Suspension and Resetting of Ca^2+^ and Transcriptional Rhythms in the Suprachiasmatic Nucleus Neurons

**DOI:** 10.1101/2022.09.18.508357

**Authors:** Ryosuke Enoki, Naohiro Kon, Kimiko Shimizu, Kenta Kobayashi, Yoshifumi Yamaguchi, Nemoto Tomomi

## Abstract

Mammalian circadian rhythms are coordinated by the master clock located in the hypothalamic suprachiasmatic nucleus (SCN). Under severe environmental conditions, such as during the harsh winter season for food, certain mammalian species reduce their basal metabolism and thermogenesis, thereby undergoing torpor, a controlled state of hypothermia, which naturally returns to the normothermic state. A long-lasting debate focused on whether the SCN with a temperature-compensated clock remains functional during hypothermia. However, so far, no direct and quantitative evidence has been reported of temperature sensitivity in living SCN neurons.

In this study, we performed dual-color fluorescence imaging of clock gene transcriptions and intracellular Ca^2+^ in mouse SCN neurons, using slices at various temperatures. We demonstrated that the *Bmal1* transcription and Ca^2+^ circadian rhythms persisted at 22°C–28°C, although the two rhythms underwent temporal dissociation at 22 °C. Notably, Ca^2+^, *Bmal1*, and *Per2* rhythms were suspended at 15°C, coupled with a significant Ca^2+^ increase, and all rhythms were reset by rewarming to 35°C. Upon rewarming, the Ca^2+^ rhythm showed stable oscillations immediately, whereas the *Bmal1* and *Per2* rhythms took several days to reach stable oscillations and recover their phase relationship with the Ca^2+^ rhythm.

Taken together, we concluded that SCN neurons remain functional under moderate hypothermic conditions at approximately 22°C–28°C but stop ticking time in deep hypothermia at 15°C and that the rhythms reset after deep hypothermia. These data also indicate that the stable Ca^2+^ oscillation precedes clock gene transcriptional rhythms in the SCN neurons.

## Introduction

In mammals, daily rhythms in physiological and biochemical events, as well as animal behavior are coordinated by the master circadian clock localized in the hypothalamic suprachiasmatic nucleus (SCN) in the brain (1). In individual SCN neurons, cellular circadian rhythms are thought to be generated by an autoregulatory transcriptional and translational feedback loop (TTFL) comprising the core and sub-loop, composed of clock genes *Period (Per) 1, Per2, Cryptochrome (Cry) 1, Cry2, Bmal1*, and *Clock*, along with their protein products (2). The circadian clock has the unique property of exhibiting a temperature-compensated rhythm with a nearly 24-hour periodicity. In general, a 10°C temperature increase accelerates the rate of biochemical reactions by a factor of 2 to 3 (Q_10_ = 2 to 3), while the Q_10_ of the circadian period is compensated to 0.8 to 1.2. This phenomenon is referred to as “temperature compensation,” an essential intrinsic circadian clock property. In addition, the TTFL is reportedly regulated by intracellular Ca^2+^ (3). A recent study described cold-responsive Ca^2+^ signaling for temperature compensation (4). Moreover, we have previously demonstrated Ca^2+^ oscillations in SCN neurons with non-functional TTFL of *Cry1/2*-deficient mice (5). This finding suggests that Ca^2+^ oscillation itself is a time-keeping mechanism that oscillates autonomously even in the absence of the TTFL.

Various mammalian species use evolutionally developed controllable hypothermia, torpor and hibernation, by lowering thermogenesis and basal metabolism to survive during the harsh winter season for food (6–8). Mammalian hibernators, such as squirrels and hamsters, undergo deep hypothermia, during which their core body temperature reaches a comparable to ambient temperature. This deep hypothermic state is maintained for several hours or days, or over a week in some species, interrupted by rewarming-related arousal through non-shivering and shivering thermogenesis. Mice enter a relatively shallow form of hypothermia, daily torpor, that reduces their energy expenditure to survive when food is unavailable at any time of the year (9). During this moderate hypothermia, the core body temperature drops to around 20-25°C and rises back to the normothermic state after several hours.

The entry of a hypothermic state, both in torpor and hibernation, is reportedly time-dependent with a circadian clock-regulated timing (8, 10, 11), although several counterarguments exist as well (12). In addition, the physiological rhythms display arrhythmia after post-hibernation (13), suggesting that the time-keeping system is affected by rewarming from low temperatures. Multiple studies have addressed the fundamental question of whether circadian rhythms continue to function during hypothermia, although in general, most reports indicate that it does not with body temperatures around 10°C (14–17). However, several studies report the presence of circadian rhythm at body temperatures above 10°C (18, 19). Since these studies were conducted in different animal species and experimental environments, continuous analysis of circadian rhythms in identical tissues under a temperature-controlled environment would be critical.

We have previously established a time-lapse fluorescence imaging approach in mouse SCN slices and discovered the cellular and network mechanisms of the circadian Ca^2+^rhythm in SCN neurons (5, 20–23). In this study, we extended our previous approach to optical imaging under a controllable temperature environment and performed simultaneous dual-color fluorescence imaging of the clock genes and intracellular Ca^2+^ rhythms in mouse SCN slices. We observed that the clock gene and Ca^2+^ circadian rhythms are maintained in the cold at 22–28°C, but suspend at approximately 15°C. Rewarming immediately resets the circadian Ca^2+^ rhythms, whereas the *Bmal1* and *Per2* rhythms gradually reestablish the phase relation relative to the Ca^2+^ rhythm. Our data demonstrate that stable Ca^2+^ oscillation precedes clock gene transcriptional rhythms in the SCN neurons.

## Results

### Low-temperature time-lapse fluorescence imaging in SCN neurons

In order to continuously monitor the circadian rhythms in the SCN neurons in a temperature-controlled environment, we constructed a time-lapse fluorescence imaging system composed of an epifluorescence microscope placed inside a thermo-regulated chamber (Fig. 1A and B, see the Materials and Methods for details). The surface of the culture medium was covered with noncytotoxic mineral oil, and the culture dish was sealed with an O_2_-permeable filter, enabling us to monitor the SCN slices without temperature difference-related water condensation and evaporation during the long day recordings (Fig. 1C). We used the yellow fluorescence protein reporter Venus under the control of a *Bmal1* promoter (Bmal1-forward-intron336-Venus-NLS-D2; *Bma1-nls-Venus*), and a red-color Ca^2+^ probe under a neuron-specific promoter (Syn1-nes-jRGECO1; nes-jRGECO1a) for visualization. We prepared mouse SCN slices and expressed these fluorescence probes by the adeno-associated virus (AAV) (Fig. 1D). To quantitatively evaluate the periodicity and continuity of the circadian rhythm, unless otherwise stated, the recording period was set for 4 days under various temperature conditions, and the total recording time was set for 12 days. We chilled the SCN slices under low temperatures, followed by rewarming to 35°C (Fig. 1E). Based on the previously reported classification of hypothermia (24), we defined 15°C, 22–28°C, and 10°C as deep, moderate, and profound hypothermia, respectively, in this study. Fig. 1F presents the typical expression patterns of nes-jRGECO1a and *Bmal1-nls-Venus* in the SCN slices. The high-resolution confocal images of the SCN slices (Fig. 1G), together with our previous studies on the Ca^2+^ probe expression patterns (21, 25), indicate that nes-jRGECO1a and *Bmal1-nls-Venus* are exclusively expressed in the SCN neuronal cytoplasm and nucleus, respectively.

**Figure 1.**
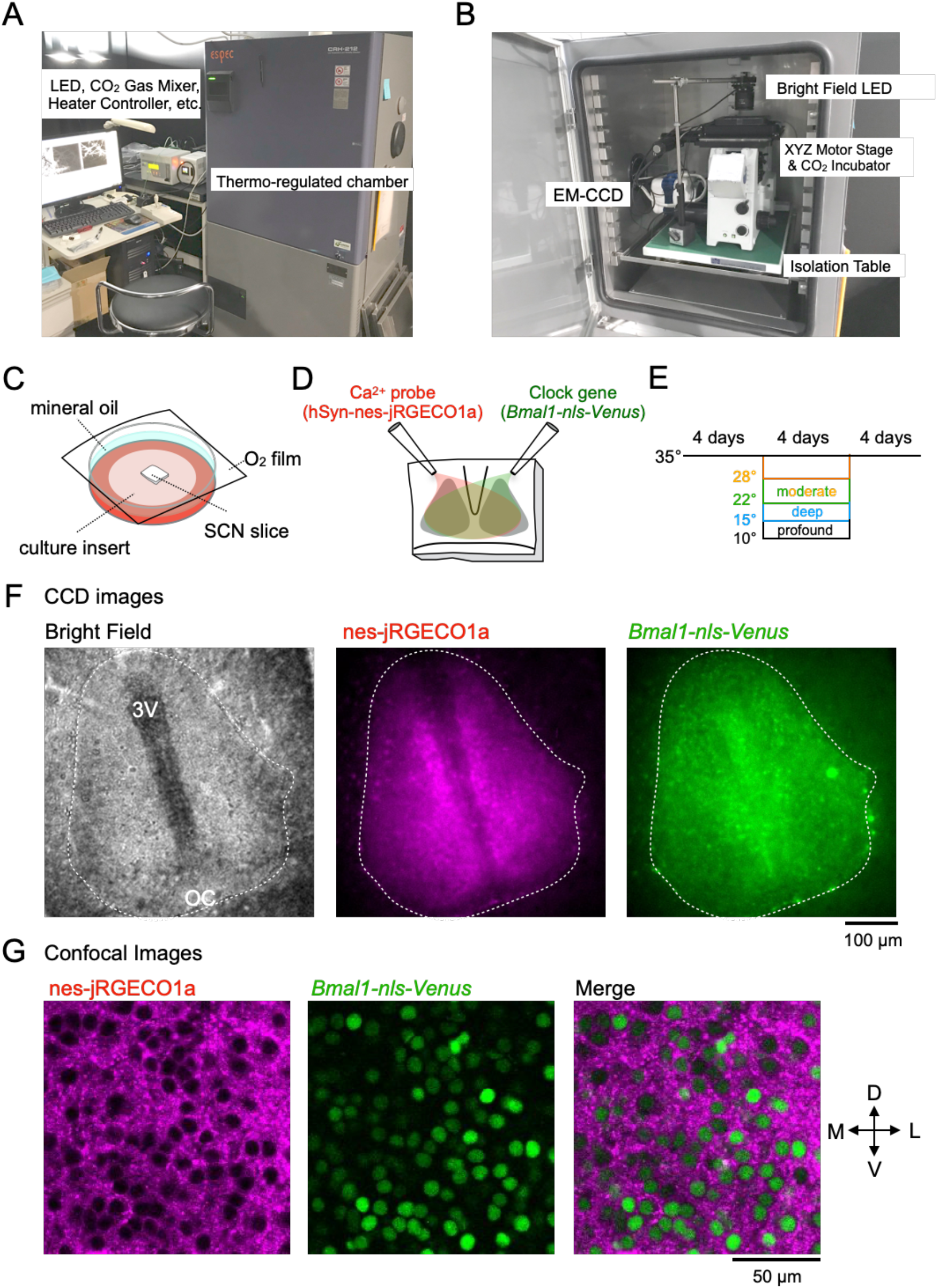
Low-temperature time-lapse fluorescence imaging in mouse SCN slices. (A) External view of the low-temperature imaging system. (B) Interior view of the microscopic platform in the thermo-regulated chamber. (C) Slice culture in the glass-bottom dish. The surface of the culture medium is covered with mineral oil, and the culture dish is sealed with an O_2_-permeable filter. (D) adeno-associated virus (AAV) infection in SCN slices. An aliquot of the AAV harboring Syn1-nes-jRGECO1a (nes-jRGECO1a) and AAV-Bmal-forward-intron336-Venus-NLS-D2 (*Bmal1-nls-Venus*) are inoculated onto the surface of the SCN slices. (E) Schedule for time-lapse imaging. After 4 days recording of the Ca^2+^ and *Bmal1* rhythms at 35°C, the SCN slices are chilled at various low temperatures for 4 days, followed by rewarming to 35°C for 4 days. In this study, we defined 15°C, 22°C and 28°C, and 10°C as deep, moderate, and profound hypothermia, respectively. (F) Representative images of mouse SCN slices expressing nes-jRGECO1a and *Bmal1-nls-Venus*. Bright-field (left), nes-jRGECO1a (center), *Bmal1-nls-Venus* (right). (G) Confocal images of nes-jRGECO1a (left), *Bmal1-nls-Venus* (center), and merged images on the left side of the SCN slices. The center region of the left SCN is depicted. 3V: the third ventricle, OC: optic chiasma.

### Ca^2+^ and *Bmal1* rhythms under cold temperatures

We monitored the Ca^2+^ and *Bmal1* signals in the SCN slices at various temperatures (35°C, 28°C, 22°C, 15°C, and 10°C) and assembled on figures the representative traces (Fig. 2), the peak phase plots of the rhythms (Fig. 3), the statistical comparison of the period and amplitude (Fig. 4), and the baseline level (Fig. 5).

**Figure 2.**
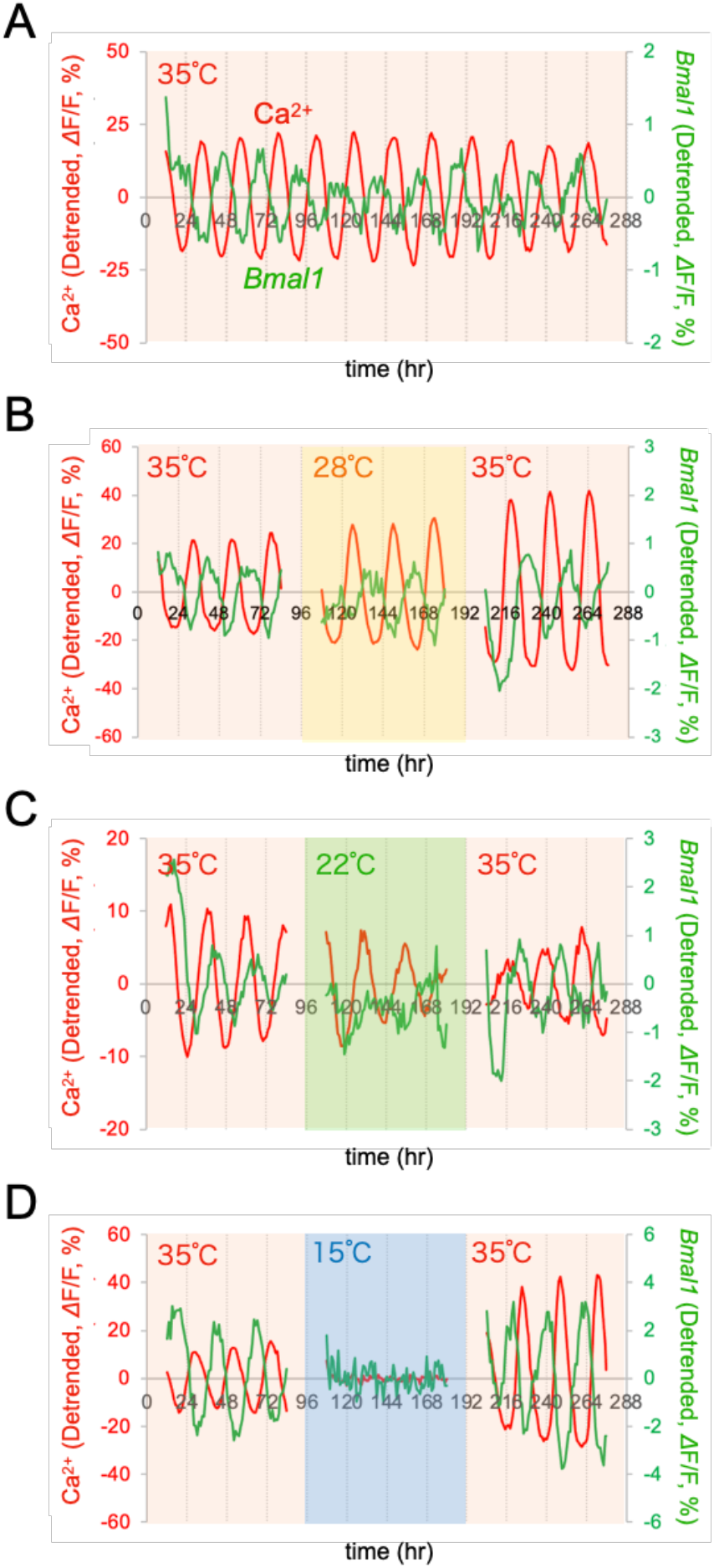
Ca^2+^ and *Bmal1* rhythms at various temperatures. Representative traces of Ca^2+^ and *Bmal1* rhythms. After recording Ca^2+^ and *Bmal1* rhythms for 4 days at 35°C, mouse SCN slices were exposed to various temperatures for 4 days (top to bottom panels: 35°C, 28°C, 22°C, and 15°C), followed by rewarming to 35°C. All traces are mean signals in the SCN region. Time is depicted after the start of the recording.

**Figure 3.**
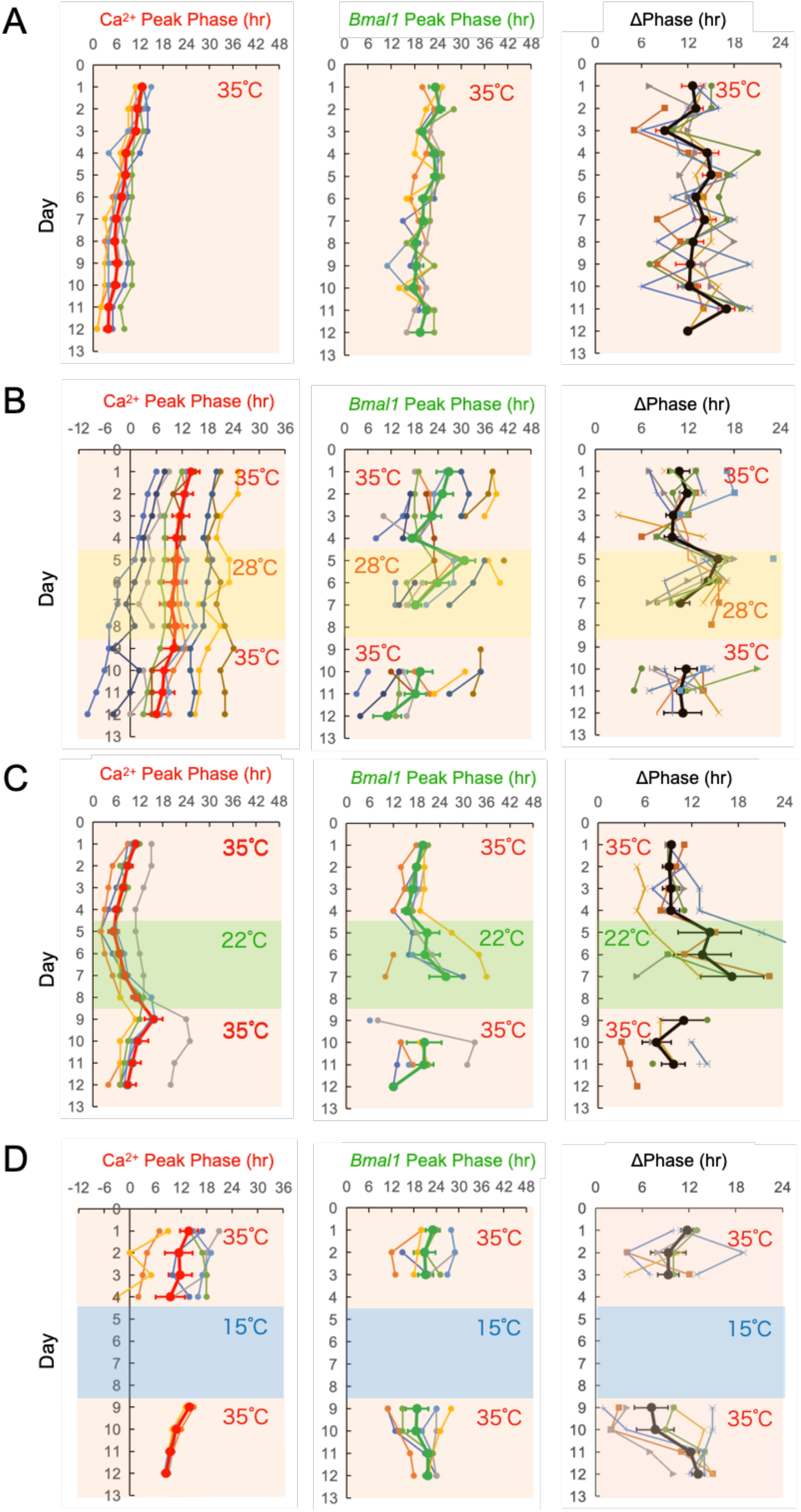
The plots of Ca^2+^ and *Bmal1* rhythms and phase difference in SCN neurons. Ca^2+^ and *Bmal1* rhythms were recorded at 35°C for 4 days, then the SCN slices were exposed to various temperatures for 4 days, followed by rewarming to 35°C for another 4 days. Ca^2+^ rhythm (left), *Bmal1* rhythm (center), and the phase difference (right). Time is depicted after the start of the recording. The data are presented as the mean ± SEM.

**Figure 4.**
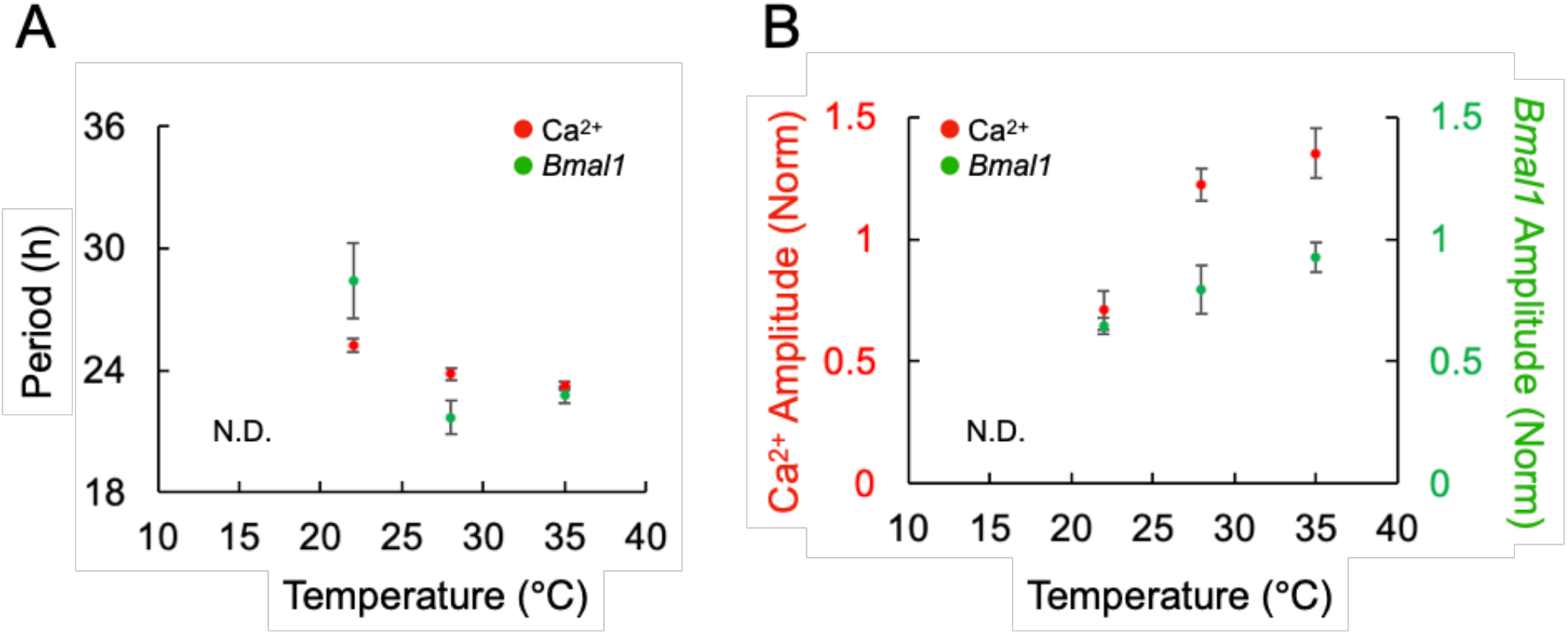
Summary of Ca^2+^ and *Bmal1* periods and amplitudes at various temperatures. Graphs represent the relationship between period (A) or amplitude (B) vs various temperatures (35°C, 28°C, and 22°C). Rhythms were not detectable at 15°C (N.D.). The amplitudes were normalized (Norm) by values before temperature change. The data are presented as the mean ± SEM.

**Figure 5.**
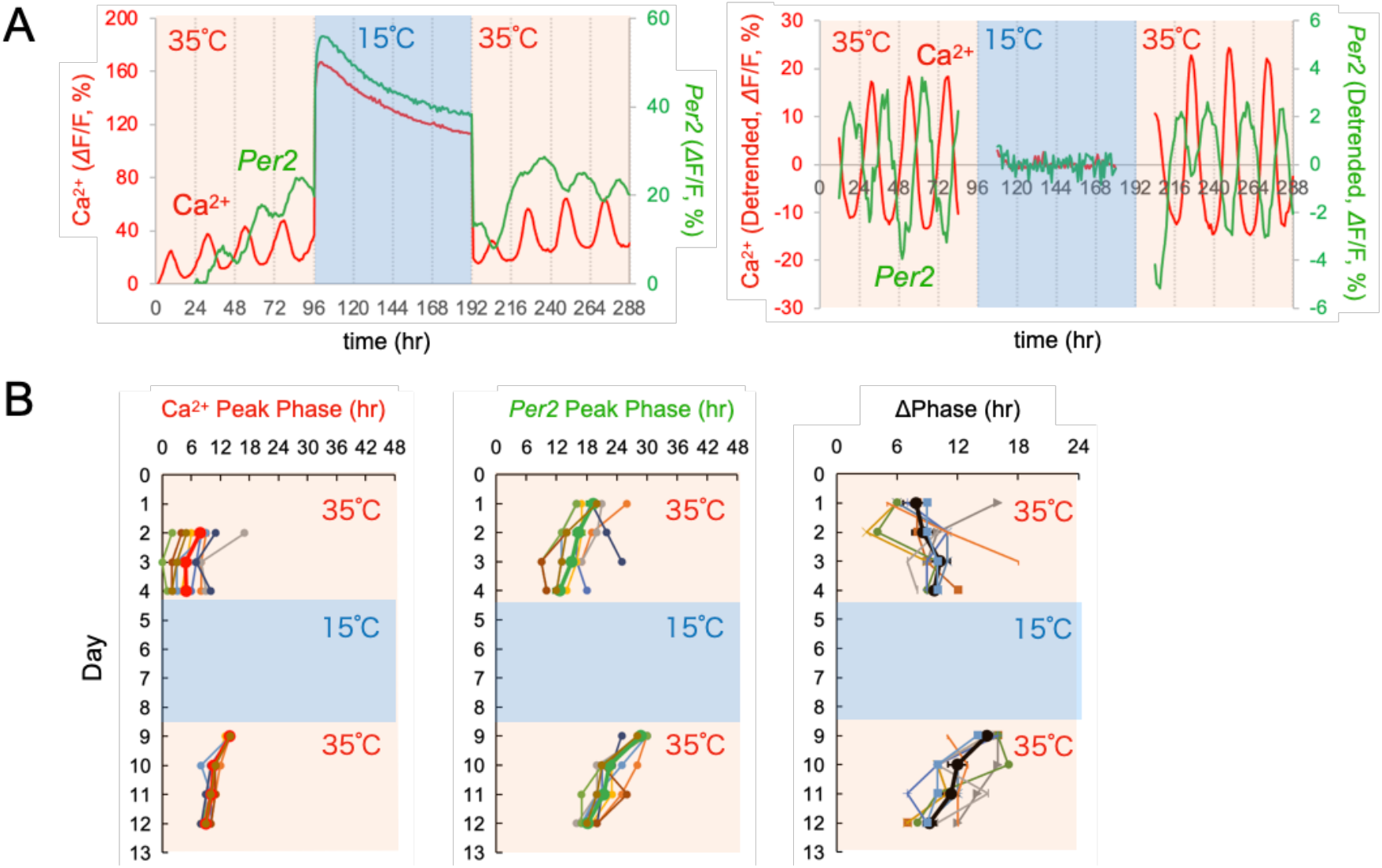
Ca^2+^ and *Per2* rhythms exposed to 15°C. (A) Representative traces of the Ca^2+^ and *Per2* rhythms. After recording Ca^2+^ and *Per2* rhythms for 4 days at 35°C, SCN slices were exposed to 15°C for 4 days, followed by rewarming to 35°C. All traces are average signals in the SCN region. (B) The plots of the peak phases of the Ca^2+^ rhythm (left), *Per2* rhythm (center), and phase difference (right). Time is depicted after the start of the recording. The data are presented as the mean ± SEM.

First, to assess the stability and continuity of the circadian rhythms under our experimental conditions, we measured Ca^2+^ and *Bmal1* signals in the SCN slices for 12 days at 35°C (Fig. 2A). We detected stable Ca^2+^ and *Bmal1* rhythms with periods slightly shorter than 24 h (Ca^2+^: 23.20 ± 0.1 h, *Bmal1*: 23.58± 0.12 h) and a nearly anti-phasic relation (13.17 ± 0.56 h) (n = 6 slices), the value of which was consistent with the results of a previous study (26).

The SCN slices were then exposed to a cold temperature of 28°C for 4 days, followed by rewarming to 35°C for another 4 days (Fig. 2B). Such low-temperature mimics moderate hypothermia present during fasting-induced torpor in mice (9) and induced hypothermia by the activation of a discrete subset of neurons in the hypothalamus (27, 28). Multiple circadian peaks could be detected in this cold (Fig. 2B), indicating the persistent rhythmicity of both Ca^2+^ and *Bmal1* (Fig. 2B). After rewarming to 35°C, both the Ca^2+^ and *Bmal1* rhythms were maintained with an anti-phase relationship. We then further chilled the SCN slices at 22°C. We observed that the Ca^2+^ and *Bmal1* rhythms persisted even under such cold conditions (Fig. 2C). After rewarming to 35°C, both the Ca^2+^ and *Bmal1* rhythms were maintained with an anti-phase relationship.

Next, the SCN slices were chilled at 15°C, mimicking deep hypothermia (Fig. 2D). Notably, neither Ca^2+^ nor *Bmal1* rhythms could be detected at this cold temperature (Fig. 2D right). After rewarming to 35°C, both the Ca^2+^ and *Bmal1* rhythms appeared and continued. Taken together, these results indicate that the circadian rhythms suspend at temperatures as cold as 15°C and that the critical point for the circadian rhythm suspension is around 15–22°C.

Under physiological conditions in mice, the core body temperature is not expected to drop below 15–22°C, although it does indeed during controlled anesthesia (29, 30). We did not detect signs of tissue damage at 15–35°C (Fig. S1A–D). However, at 10°C, especially after rewarming to 35°C, we detected severe tissue damage and shrinkage (Fig. S1E). These results indicate that the SCN in mice cannot tolerate exposure to cold temperatures below 10°C, we thus excluded these data from further analysis.

### Statistical evaluation of Ca^2+^ and *Bmal1* rhythms under cold temperatures

At a constant temperature of 35°C, the Ca^2+^ and *Bmal1* rhythms with an anti-phasic relationship could be detected for 12 days in the SCN neurons (Fig. 3A) (n = 6 slices). The Ca^2+^ and *Bmal1* rhythm periods were shorter than 24 hours, remaining stable during the long recording for 12 days (Ca^2+^ rhythm: 22.61 ± 0.50, 23.25 ± 0.19, and 23.58 ± 0.14 h on Days 1–4, 5–8, and 9–12, respectively; *Bmal1* rhythm: 23.25 ± 0.48, 22.79 ± 0.37, and 24.81 ± 0.58 h on Days 1–4, 5–8, and 9–12, respectively) (Fig. S2A1). The Ca^2+^ and *Bmal1* rhythm amplitudes were also stable over the long recording (Ca^2+^ rhythm: 23.30% ± 4.74%, 29.84%± 4.78%, and 29.91%± 5.16% on Days 1–4, 5–8, and 9–12, respectively; *Bmal1* rhythm: 1.52% ± 0.33%, 1.35% ± 0.27%, and 1.45% ± 0.37% on Days 1–4, 5–8, and 9–12, respectively) (Fig. S2A2).

Under cold temperatures of 28°C, the Ca^2+^ rhythm phase remained stable, whereas the *Bmal1* rhythm was transiently delayed (Fig. 3B, center), and the phase difference between the Ca^2+^ and *Bmal1* rhythms widened (Day 5 vs Day 4, *p* = 0.0482). After this transient delay, the *Bmal1* rhythm phase advanced and the phase difference returned to the prechilling level on Day 7. At 28°C, the Ca^2+^ rhythm period slightly lengthened (Fig. S2B1) (22.85 ± 0.14, 23.88 ± 0.29, and 22.84 ± 0.21 h at 35°C, 28°C, and after rewarming, respectively; *p* = 0.006) (n = 10 slices), whereas the *Bmal1* rhythm did not (23.90 ± 1.07, 21.67 ± 0.82, and 22.85 ± 1.34 h at 35°C, 28°C, and after rewarming, respectively). Our statistical analysis showed that the *Bmal1* rhythm amplitudes became significantly smaller (1.53% ± 0.16%, 1.08% ± 0.07%, and 1.74% ± 0.19% at 35°C, 28°C, and after rewarming, respectively; *p*=0.034), although it was not the case for Ca^2+^ rhythms (31.91% ± 3.84%, 39.93% ± 5.70%, and 43.14% ± 9.52% at 35°C, 28°C, and after rewarming, respectively) (Fig. S2B2). These results show that both the Ca^2+^ and *Bmal1* circadian rhythms persist at as moderate cold temperatures as 28°C and that the Ca^2+^ rhythm remains stable, whereas the *Bmal1* rhythm is more sensitive to temperature changes.

At the cold temperature of 22°C, the peak phases of the Ca^2+^ and *Bmal1* rhythms were delayed (Fig. 3C, left and center) (n = 6 slices) and the Ca^2+^ and *Bmal1* rhythm phase differences gradually widened during this cold exposure (Fig. 3C, right). The periods of the two rhythms significantly lengthened (Ca^2+^ rhythm: 22.39 ± 0.10, 25.25 ± 0.33, and 22.14 ± 0.26 h at 35°C, 28°C, and after rewarming, respectively, *p* = 0.0006; *Bmal1* rhythm: 22.50 ± 0.36, 28.40 ± 1.83, and 24.88 ± 0.97 h at 35°C, 28°C, and after rewarming, respectively, *p* = 0.0342) (Fig. S2C1). Notably, the *Bmal1* period was significantly larger in Ca^2+^ rhythms (*p* = 0.0482). After rewarming, the period was returned to that of the pre-chilling levels (Fig. S2C1). The Ca^2+^ and *Bmal1* rhythm amplitudes were significantly attenuated (Ca^2+^rhythm: 34.87% ± 5.31%, 25.17% ± 5.25%, and 26.56% ± 6.79% at 35°C, 28°C, and after rewarming, respectively, *p* = 0.0388; *Bmal1* rhythm: 1.96% ± 0.22%, 1.27% ± 0.17%, and 2.04% ± 0.23% at 35°C, 28°C, and rewarming, respectively, *p* = 0.0006) (Fig. S2C2). These results showed that both the Ca^2+^ and *Bmal1* rhythms persist at the temperature of 22°C and these two rhythms transiently dissociate under such cold conditions.

Upon cold exposure at 15°C, we could not detect any Ca^2+^ and *Bmal1* rhythm (Fig. 3D). Notably, upon rewarming, the Ca^2+^ rhythms were restarted at a certain phase, while *Bmal1* reached a stable phase after a longer transient state (Fig. 3D, left and center). The Ca^2+^ rhythm peak phases could be detected at 14.00 ± 0.26 h after rewarming (n = 6), whereas those of *Bmal1* were detected widely on Day 1 after rewarming (SD = 7.94 h).

Fig. 4 summarizes the temperature-dependent changes in the Ca^2+^ and *Bmal1* rhythm periods and amplitudes. The Ca^2+^ rhythm periods were relatively constant compared to those of the *Bmal1* rhythms (Fig. 4A). The amplitude of both rhythms decreased monotonically (Fig. 4B). The critical temperature for the rhythm suspension was at approximately 15°C and below 22°C, respectively.

### Cold exposure increases the Ca^2+^ and *Bmal1* baseline levels

Cold exposure at 22°C–28°C increased the Ca^2+^ rhythm baseline levels (Fig. S3A). Notably, gradual fluorescence intensity increases could be detected during the long recordings, typically occurring in AAV-transfected SCN neurons (20, 21, 31) (Fig. S3). Fig. S3B presents the Ca^2+^ baseline level comparisons on Days 1 and 4 (Day 1: 28.69% ± 9.17%, 48.21% ± 14.65%, and 68.44% ± 11.00% at 35°C, 28°C, and 22°C, respectively; Day 4: 56.68% ± 13.30%, 96.57% ± 23.63%, and 71.94% ± 11.65% at 35°C, 28°C, and 22°C, respectively). At 15°C, the Ca^2+^ baseline level increased rapidly and reached significantly higher levels on Day 1 (196.70% ± 12.31%, *p* < 0.0001) and Day 4 (152.22% ± 12.58%, *p* = 0.0104). After rewarming to 35°C, the baseline levels returned quickly to lower values (Fig. S3A). In contrast, the *Bmal1* baseline level gradually increased during the 22°C–28°C exposure (13.37% ± 0.82%, 72.49% ± 7.77%, and 107.10% ± 24.66% at 35°C, 28°C, and 22°C, respectively, *p* = 0.0037 for 28°C, *p* < 0.0001 for 22°C) (Fig. S3C and D). In the case of 15°C cold exposure, the increase in the baseline level of the *Bmal1* rhythm was smaller and flattened (25.39% ± 2.94%). These results show that cold temperature increases the Ca^2+^ and *Bmal1* baseline levels and the rhythms suspend at 15°C accompanied by a significant Ca^2+^ increase.

### Rewarming differentially restarts the *Bmal1* and *Per2* circadian rhythms

Next, we compared the cold responses of two transcriptional rhythms, *Bmal1* and *Per2*. The *Bmal1* transcription depends on RORE in the upstream region of *Bmal1*, whereas *Per2* transcription is regulated through the cAMP responding element (CRE) or E-box by CREB or CLOCK (1,2). Because CREB and CLOCK are regulated by Ca^2+^-CaMKII (32), *Per2* is directly regulated by Ca^2+^ signaling.

We expressed Venus under *Per2* promoter (*pAAV-Per2-intron2-Venus-NLS-D2; Per2-nls-Venus*) together with nes-jRGECO1a in the SCN slices. Similar to in the case of *Bmal1*, we confirmed that *Per2-nls-Venus* was expressed exclusively in the nuclei of the SCN neurons (Fig. S6). After confirming that the Ca^2+^ and *Per2* rhythms were stable with a phase difference (7.88 ± 1.25 h), the SCN slices were cooled at 15°C for 4 days, followed by rewarming to 35°C for another 4 days (Fig. 5). We observed that, as in the case of *Bmal1*, the Ca^2+^ and *Per2* rhythms were not detectable at 15°C, accompanied by a significant baseline level increase (Fig. 5A). Notably, unlike in the case of *Bmal1*, both the Ca^2+^ and *Per2* rhythms were restarted quickly on Day 1 after rewarming (Fig. 5B, left and center). The peak phases of the Ca^2+^ and *Per2* rhythms were detected at 13.89 ± 0.11 h and 4.78 ± 0.55 h after rewarming, respectively (n = 9). In particular, the Ca^2+^ and *Per2* rhythm phase differences were large on Day 1 after rewarming (14.89 ± 0.45 h), but gradually recovered toward the original phase difference after a transient state for 4 days (9.22 ± 0.62 h) (Fig. 5, right).

Fig. 6 presents the Rayleigh plot analysis, showing that the Ca^2+^ rhythm restarted on Day 1 and the *Bmal1* rhythm gradually converged to a similar phase on Day 4, re-establishing the anti-phase relationship (Fig. 7A). However, the *Per2* rhythm restarted on Day 1, although with a larger phase difference relative to that of the Ca^2+^ rhythms. The Ca^2+^ and *Per2* phase differences were gradually re-established after a transient period of 4 days (Fig. 7B). These results suggest that the Ca^2+^ rhythm differentially sets the phase of the *Bmal1* and *Per2* transcriptional rhythms, and the extent of which is greater in *Per2* than *Bmal1*.

**Figure 6.**
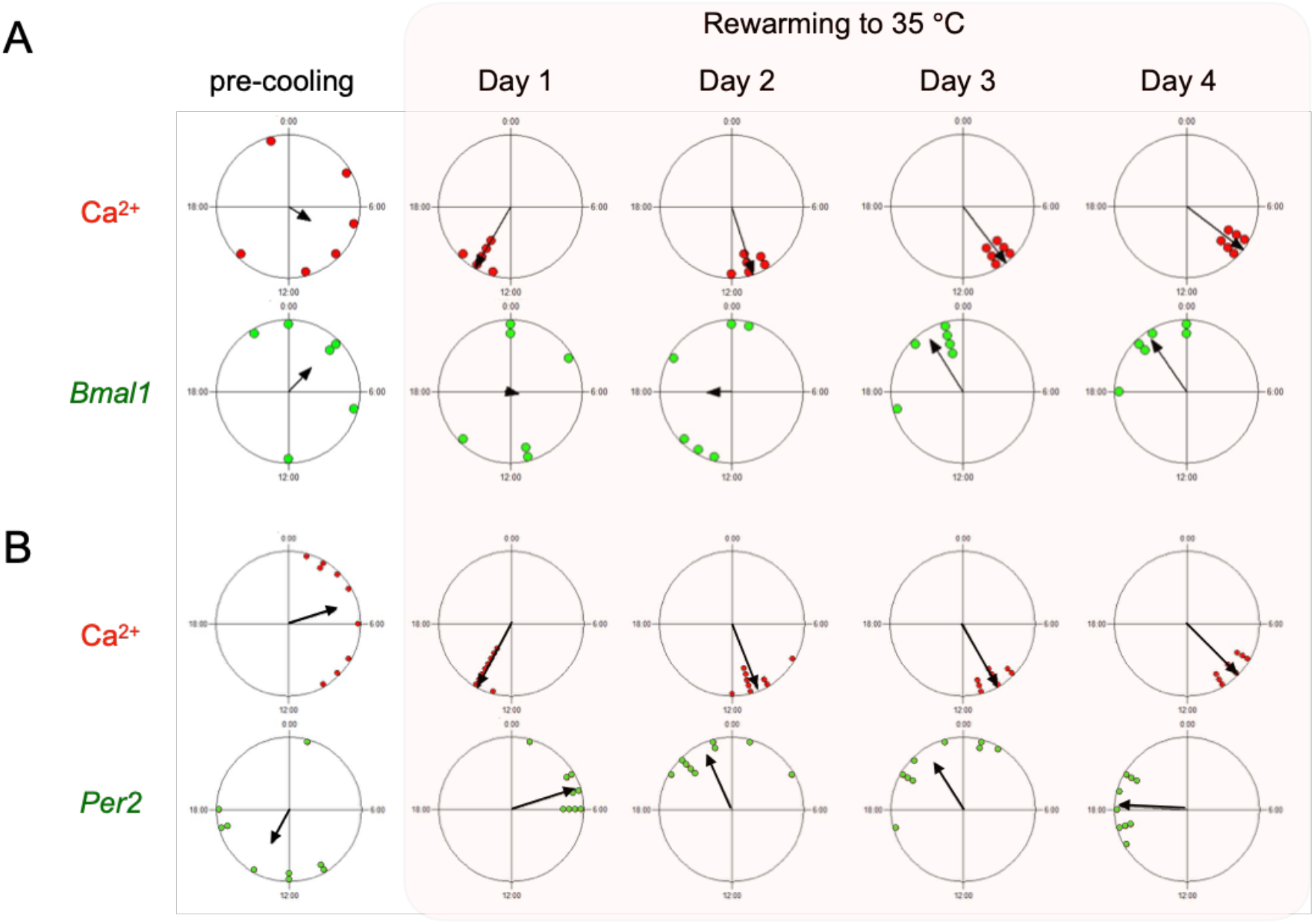
Rayleigh plots of Ca^2+^, *Bmal1*, and *Per2* rhythms before cooling and after rewarming. Rayleigh plots of the peak phase of the Ca^2+^ and *Bmal1* rhythms (A) and the Ca^2+^ and *Per2* rhythms (B). SCN slices were exposed under 15°C for 4 days, and the peak phases before cold exposure (pre-cooling) and after rewarming were calculated with respect to the start and end of cold exposure. The small circles on the Rayleigh circles indicate the phase of individual SCN slices, and the arrow lines within the circles indicate the average phase.

**Figure 7.**
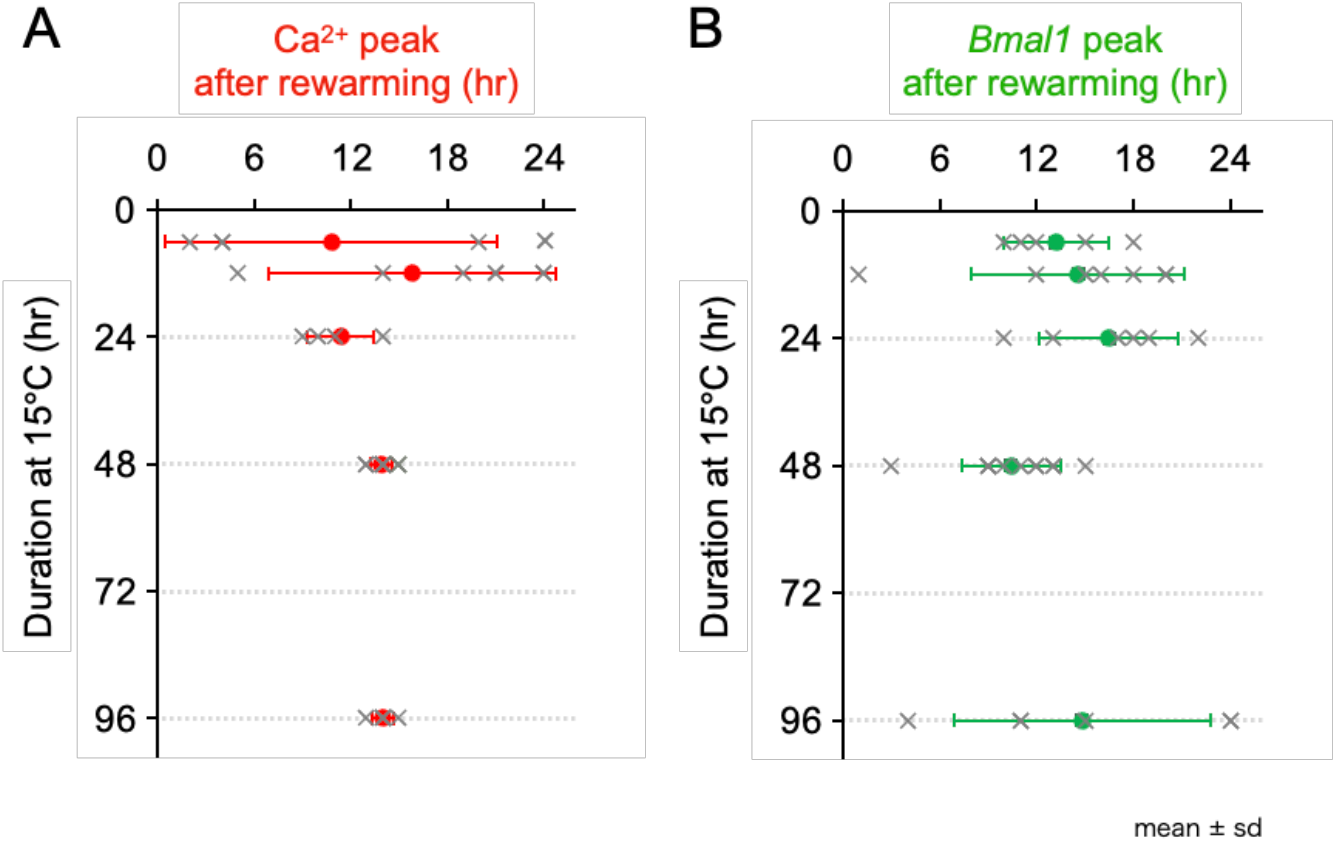
Relationship between cold exposure duration and peak phase after rewarming in SCN neurons. The 1^st^ peak phases of Ca^2+^ (A) and *Bmal1* (B) after rewarming. The duration of cold exposure at 15°C systematically varied from 6, 12, 24, 48, and 96 h. Individual data are represented by gray color x and mean ± SD by red/green colors.

### Time required to rhythm reset

Finally, we examined the time necessary to reset the Ca^2+^ rhythm after rewarming. The SCN slices were exposed to 15°C and the duration of cold exposure was systematically varied for 6–48 h. Fig. S4 and Fig. S5 present the representative Ca^2+^ and *Bmal1* rhythm traces and phase plots, respectively. Short cold exposures (6–24 h) delayed the phase but did not reset the rhythms (Fig. S4A–C and Fig. S5A–C). Notably, the rewarming from a 48-h cold exposure reset the Ca^2+^ rhythms on Day 1. In contrast, the *Bmal1* rhythm phases were widespread on Day 1 (SD = 9.48 h), but they gradually converged into an identical phase on Day 4 (SD = 1.63 h) (Fig. S4D and S5D) and re-established the anti-phase relationship relative to the Ca^2+^ rhythms (13.00 ± 1.22 h).

Fig. 7 summarizes the relationship between the duration of cold exposure and the 1^st^ peak phase of the rhythms after rewarming. After 6–24 h of cold exposure at 15°C, the Ca^2+^rhythms were not reset (6, 12, and 24-h exposures: 10.80 ± 10.35, 15.85 ± 8.94, and 11.40 ± 2.07 h, respectively), but a 48-h cold exposure fully reset the Ca^2+^ rhythms (13.92 ± 0.67 h). In contrast, the *Bmal1* rhythms were not reset by any of the cold exposures (6, 12, 24, and 48-h exposures: 13.20 ± 3.27, 14.57 ± 6.63, 16.50 ± 4.32, and 10.42 ± 3.06 h, respectively). These results suggest that a cold exposure over 48 hours would be required for the phase resetting of the Ca^2+^ rhythms.

## Discussion

Whether the clock gene transcriptional oscillations in the SCN neurons continue under hypothermia, specifically in the case of hibernation and daily torpor, has long been debated (15, 16). A major reason for this question remaining unresolved is that brain samples were obtained from different individual animals in order to estimate the temporal changes in clock genes and their protein expressions (15, 16). This was based on the assumption that the circadian rhythm phase during hypothermia is identical in all animals. However, the rhythm analysis accuracy would be significantly reduced if individual differences in animals were present. To solve these questions, it was necessary to evaluate circadian rhythms from identical living SCN tissue samples held in a temperature-controlled environment. Moreover, a growing body of evidence supports the existence of a non-transcriptional oscillator without key clock genes (33, 34), raising the possibility that the expression of the clock genes might not be an appropriate index for assessing circadian rhythms under hypothermic conditions including torpor and hibernation. We have recently demonstrated that Ca^2+^ signaling compensates for the slow transcriptional oscillation speed at low temperatures (4). Furthermore, we reported circadian Ca^2+^ rhythm persistence in SCN neurons lacking key clock genes *Cry 1/2* (31). These results suggest that not only are Ca^2+^ rhythms important for TTFL but they themselves are important for self-sustained, temperature-compensated oscillation. In this study, we attempted to elucidate the mechanism of SCN rhythmicity by simultaneously measuring clock genes as well as Ca^2+^under cold temperature conditions.

### Circadian rhythms under hypothermia

We discovered that the circadian rhythms of Ca^2+^, *Bmal1*, and *Per2* in the mouse SCN neurons suspended at a cold temperature of 15°C. These results are consistent with those of previous reports using immunostaining or *in situ* hybridization in the brain samples of hibernating animals (15, 16). During such deep hypothermia, clock gene and protein expressions were up-regulated and suspended, and a neural activity marker was up-regulated in the SCN. Interestingly, similar to the present results, the cyanobacterial circadian clock is lost at low temperatures around 19°C, and the self-sustained rhythm of cyanobacterial KaiC phosphorylation transformed to damped oscillations, as predicted by the Hopf bifurcation theory (35). Although the key molecules responsible for circadian oscillations differ between mammalian cells and cyanobacteria, the rhythm suspend at similarly low temperatures is intriguing from the aspect of clock function evolution.

In this study, the baseline level of Ca^2+^, *Bmal1*, and *Per2* was up-regulated at low temperatures. Increased Ca^2+^ might activate the SCN neurons, thereby increasing clock gene and protein expressions. Ca^2+^ increase at as cold temperatures as 15°C was significant and the rise and fall times of Ca^2+^ changes were rapid, i.e., within 1–2 hours. Cooling reportedly increases Ca^2+^ and tension in skeletal muscle (36), which is mediated by Ca^2+^release from the internal stores. The origin of the cold-induced Ca^2+^ increase remains unknown in the SCN neurons, although a similar mechanism might exist. The amplitude of the circadian Ca^2+^ rhythm reported in SCN neurons is approximately 50-100 nM (37, 38). In this study, the amplitude of the Ca^2+^ rhythm was ca. 20%–30% (Fig. S2) and the Ca^2+^increase magnitude at 15°C was 200% in SCN neurons (Fig. S3A and B), respectively. Therefore, the Ca^2+^ increase magnitude under severe cold is estimated to be approximately 1 μM. Ca^2+^ signaling at low temperatures could be a key factor for the rhythm suspension and resetting.

### Theoretical insight: the hierarchy of Ca^2+^ oscillation and TTFL

In this study, we examined the time required to reset the circadian rhythms. Our results showed that 6–24 hours of cold exposure was insufficient, but 48 hours of exposure fully reset the Ca^2+^ rhythms in the SCN neurons. Upon rewarming, the *Bmal1* phase was widely distributed on Day 1, and the *Bmal1* rhythms gradually re-established the phase relation relative to the Ca^2+^ rhythms (Fig. 6A). In comparison, the Ca^2+^ and *Per2* rhythms immediately restarted on Day 1. The *Per2* rhythm was phase-shifted gradually during the transient period and the original phase difference was restored relative to the Ca^2+^ rhythms (Fig. 6B).

These differential properties in the Ca^2+^, *Bmal1*, and *Per2* rhythms could be explained by the limit cycle model, in which the amplitude of the limit cycle becomes smaller with decreasing temperatures (Fig. 8A). The critical temperature for the rhythm suspension, a point of Hopf bifurcation, is 15°C–22°C. The baseline Ca^2+^ level increased with the decreasing temperature and the transcription rhythm suspension were accompanied by relevant Ca^2+^ increases, suggesting that Ca^2+^ signaling regulates the clock gene transcription rhythms. At temperatures as cold as 22°C, the Ca^2+^ and *Bmal1* rhythms were transiently dissociated (Fig. 4A), implying the presence of multiple oscillators in the SCN neurons.

**Figure 8.**
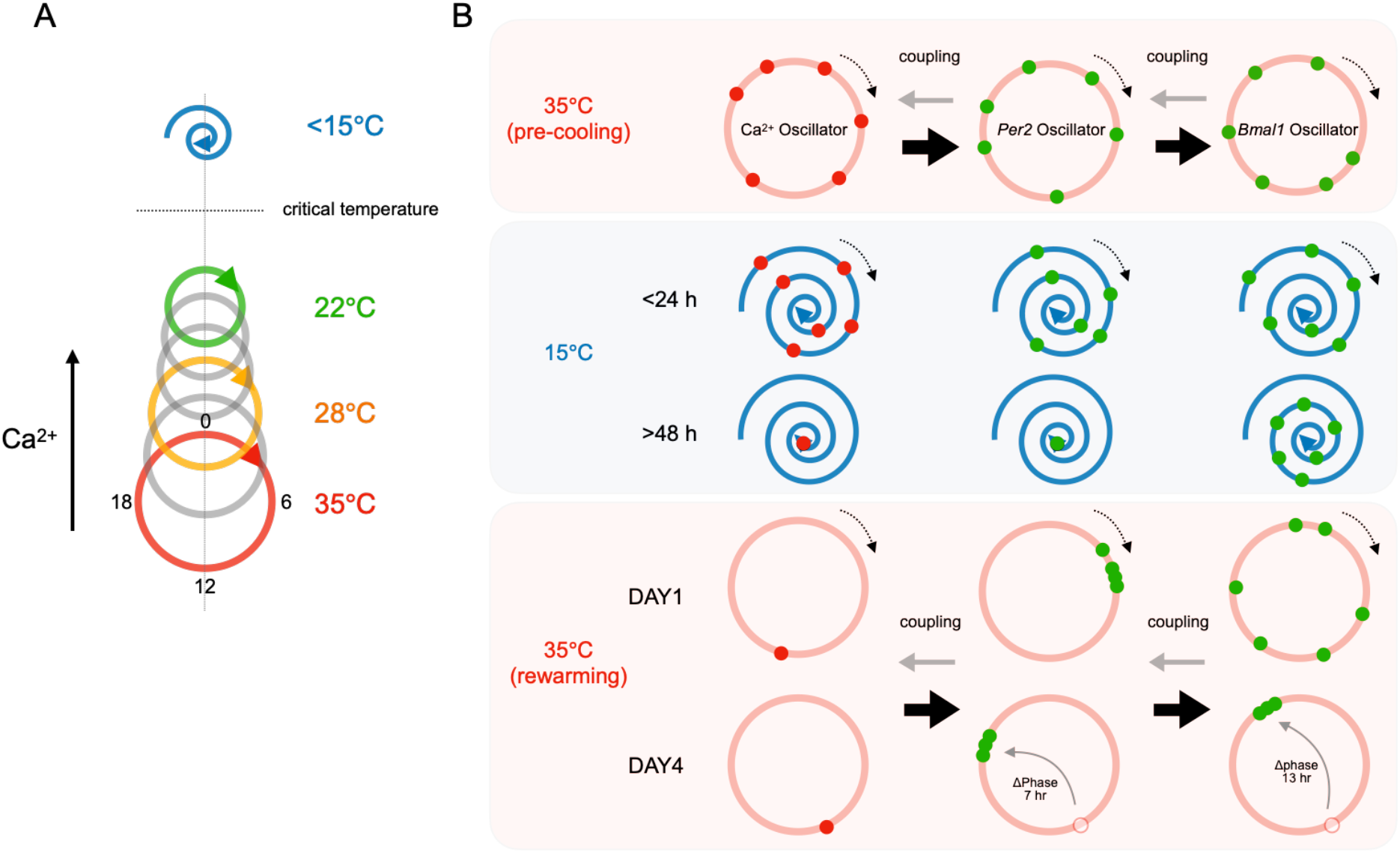
Limit cycle model of cold-responsive circadian rhythms. (A) The amplitude of the limit cycle in the Ca^2+^ and clock gene transcription (*Bmal1/Per2*) decreases with a decreasing temperature. Under severe cold at 15°C, the limit cycles are transformed into the damping oscillator. The critical temperature for the rhythm suspension is around 15°C–22°C. The Ca^2+^ level sets the size of the limit cycles. (B) Under severe cold at 15°C, the Ca^2+^ and *Per2* rhythms slow down and eventually suspend at a certain point after 48 h cold exposure, then restart from the identical phase after rewarming to 35°C. However, the *Bmal1* limit cycle damps slowly and suspends at various phases, and the phases after rewarming are variable and take several days to resynchronize to the Ca^2+^rhythms with anti-phase relation. The *Bmal1* and *Per2* phases, relative to the Ca^2+^ phase, are gradually restored during a transient period.

*Per2* constitutes the TTFL core loop, regulated by Ca^2+^ via CaMKII and CRE (1, 32). In comparison, *Bmal1* constitutes a TTFL sub-loop, not considered to be directly regulated by Ca^2+^. Taken together, we hypothesize the hierarchy of the Ca^2+^, *Per2*, and *Bmal1* oscillators. The Ca^2+^ oscillator regulates the *Per2* and *Bmal1* oscillations, probably in this order (Fig. 8B, upper row in red). Under cold temperature conditions of 15°C, the Ca^2+^ and *Per2* limit cycles transformed into damped oscillators and eventually suspend at a certain phase (Fig. 8B, middle row in blue), and restart from the identical phase after rewarming to 35°C (Fig. 8B, lower row in red). In contrast, the *Bmal1* limit cycle damps slowly and suspends at various phases, and the phases after rewarming are variable and take several days to reestablish the phase relation with the Ca^2+^ rhythms. The phase difference between Ca^2+^and *Per2* was initially beyond 12 hours, but was gradually restored to its original value of ca. 7 hours (Fig. 6). These data suggest that the Ca^2+^ oscillator leads the phase of the oscillations of *Bmal1* and *Per2*.

### Perspectives

In this study, fluorescence time-lapse imaging on mouse SCN slices in a temperature-controlled environment allowed us to investigate in detail the characteristics of circadian rhythms at low temperatures. It has been reported that the physiological rhythms of hibernators are arrhythmic in the post-hibernation (13). The present study revealed unstable transcriptional rhythms in the SCN upon rewarming from cold temperatures. These results indicate that the unstable physiological rhythms of hibernators might be due to unstable SCN oscillations. However, the *in vivo* rhythms in adult hibernators remain unknown. To address this aspect, direct and continuous measurements from the deep brain regions of freely moving hibernators would be required. In addition, the cellular and molecular mechanisms by which rhythms are arrested at low temperatures and restart upon rewarming are unknown, and these would be important questions to be addressed in future research.

## Materials and Methods

### Animal care

We obtained female mice with newborn pups (C57BL/6J) from the animal breeder (Japan SLC, Inc.). The animals were housed in our animal quarters under controlled environmental conditions (temperature: 22°C ± 2°C, humidity: 40% ± 20%, 12-h light/12-h dark, with lights on from 0800–2000 hours). The light intensity was adjusted to approximately 100–200 l× at the cage surface. The animals were fed commercial chow (Labo MR Standard; Nosan Corporation) and tap water. All animal care and experimental procedures were approved by the Institutional Animal Care and Use Committee of the National Institute of Natural Sciences and performed according to the National Institute for Physiological Sciences guidelines (Approved No. 20A017 and 20A122).

### SCN slice culture

The brains of neonatal mice (5-to-6-day-old, both male and female) were obtained in the middle of the light phase under hypothermic anesthesia, and rapidly dipped into an ice-cold balanced salt solution comprising 87 mM NaCl, 2.5 mM KCl, 7 mM MgCl_2_, 0.5 mM CaCl_2_, 1.25 mM NaH_2_PO_4_, 25 mM NaHCO_3_, 25 mM glucose, 10 mM HEPES, and 75 mM sucrose. A 200-μm coronal slice containing the mid-rostro-caudal region of the SCN was carefully prepared using a vibratome (VT 1200; Leica) and explanted onto a culture membrane (Millicell CM; pore size, 0.4 μm; Millipore) in a 35-mm Petri dish containing 1 mL of Dulbecco’s Modified Eagle Medium (DMEM) (Invitrogen) and 5% fetal bovine serum (Sigma–Aldrich). Prior to recording, the culture membrane was transferred onto a glass-bottom dish (35 mm, 3911-035; IWAKI) and the dishes were sealed with O_2_-permeable filters (membrane kit, High Sens; YSI) using silicone grease compounds (SH111; Dow Corning Toray). We applied 500 μL of Mineral Oil (Sigma–Aldrich) onto the surface of the DMEM to prevent evaporation and condensation during the long recording in the temperature-changing environment. We added 10 μM α-tocopherol (Tokyo Chemical Industry Co., Ltd.) to DMEM to prevent ferroptosis (39).

### AAV-mediated gene transfer into the SCN slices

*pAAV-Bmal-forward-intron336-Venus-NLS-D2* was generated by swapping the *Cry1* promoter of *pAAV-P(Cry1)-forward-intron336-Venus-NLS-D2* (Addgene) with a *Bmal1* promoter (40). *pAAV-Per2-intron2-Venus-NLS-D2* was generated by deleting the LoxP cassettes of *pAAV-P(Per2)-DIO-intron2-Venus-NLS-D2* (Addgene). The AAV vectors were produced by a procedure as described previously (41). AAV1-Syn1-nes-jRGECO1a was purchased from Addgene. AAV aliquots (0.8 μL) were inoculated onto the surface of the SCN slices on Days 4–5 of culture. The infected slice was cultured for a further 9–12 days before imaging. The titer of Syn1-nes-jRGECO1a, *Bmal1-nls-Venus*, and *Per2-nls-Venus* was over 1.2 × 10^13^ genome copies/mL.

### Intracellular Ca^2+^ and clock gene expression imaging

Bioluminescence imaging using luciferase reporter is a generally used standard method to observe circadian rhythms (42). However, it is based on a biochemical reaction and metabolism, such as luciferin-luciferase enzymatic reaction and the hydrolysis of ATP, the activity of which is also considered to be temperature-dependent. Therefore, we used fluorescence reporters to assess Ca^2+^ and transcription rhythmicity. The low-temperature time-lapse imaging system was composed of an EM-CCD camera (Evolve; Photometrics), an inverted microscope (IX70; Olympus), dry objectives (20 X, 0.75 NA, UPlanSApo; Olympus), a stage incubator (CU-109; Live Cell Instrument), an XYZ controller, a filter wheel (MAC6000, Ludl Electronic Products) and a LED driver (LEDD1B, Thorlabs). The microscopic system was placed in the low-temperature controller (CRH-212, ESPEC CORP). The upper part of the inverted microscope was removed to fit into the low-temperature controller. The microscopic system was controlled by the MetaMorph software (Molecular Devices).

The time-lapse wide-field imaging was conducted with an exposure time of 100 msec at 1-hour intervals. Venus and jRGECO1a were excited by cyan (480/17 nm) and yellow color (580/14 nm) with the LED light source (Spectra X Light Engine; Lumencor Inc) and the fluorescence was visualized with a dual-edge dichroic mirror (FF505/606-Di01, Semrock) and emission filters (FF01-530/43, FF02-641/75). We continuously perfused 5% CO_2_ with a gas mixer (GM-2000, Tokai-hit). Tissue and cell conditions were monitored by cultured slice bright-field images and calcium signals in individual neurons.

### Data analysis and statistics

Statistical analyses were performed using Excel (Microsoft) and Prism GraphPad (GraphPad Software). Circular plots were generated using Oriana (Kovach Computing Services). Paired or unpaired t-tests were used when comparing two dependent and independent group means. One-way ANOVA with a post-hoc Dunnett’s post-test was used to validate the temperature effects when paired multiple group data were compared.

## Acknowledgments

We are grateful to the members of the Biophotonics Laboratory, the Transformative Research Area “Hibernation Biology,” Gen Kurosawa (RIKEN iTHEMS), and Hiroshi Ito (Kyushu University) for the helpful discussions of this study. We also thank the Genetically-Encoded Neuronal Indicator and Effector Project and the Janelia Farm Research Campus of the Howard Hughes Medical Institute for sharing the jRGECO1a constructs; Maki Watanabe and Yoshiko Yamada for the excellent technical support on the experiments; Yuki Watakabe, Miwa Kawachi, and Chiemi Hyodo for their help with animal care and laboratory management. This work was supported by the Ministry of Education, Culture, Sports, Science, and Technology (MEXT)/Japan Society for the Promotion of Science KAKENHI to R.E. (20H05769, 20H03425, 22K19319), to T.N. (20H05669, 20H00523), and to Y.Y. (20H05766), Japan Agency for Medical Research and Development (AMED) to Y.Y. (22gm6310019), K.S. (21H02524), Takeda Science Foundation, the Mochida memorial foundation, Inamori Research Institute for Science, Joint Research of the ExCELLS (No, 21-205), and Cooperative Study Program (no, 21-136 and no, 22-159) of the National Institute for Physiological Sciences.

## Data and material availability

All data needed to evaluate the conclusions in the study are present in the main text and the Supplementary Materials. Additional data related to this study could be requested from the authors.

---

**Supplemental Figure 1.**
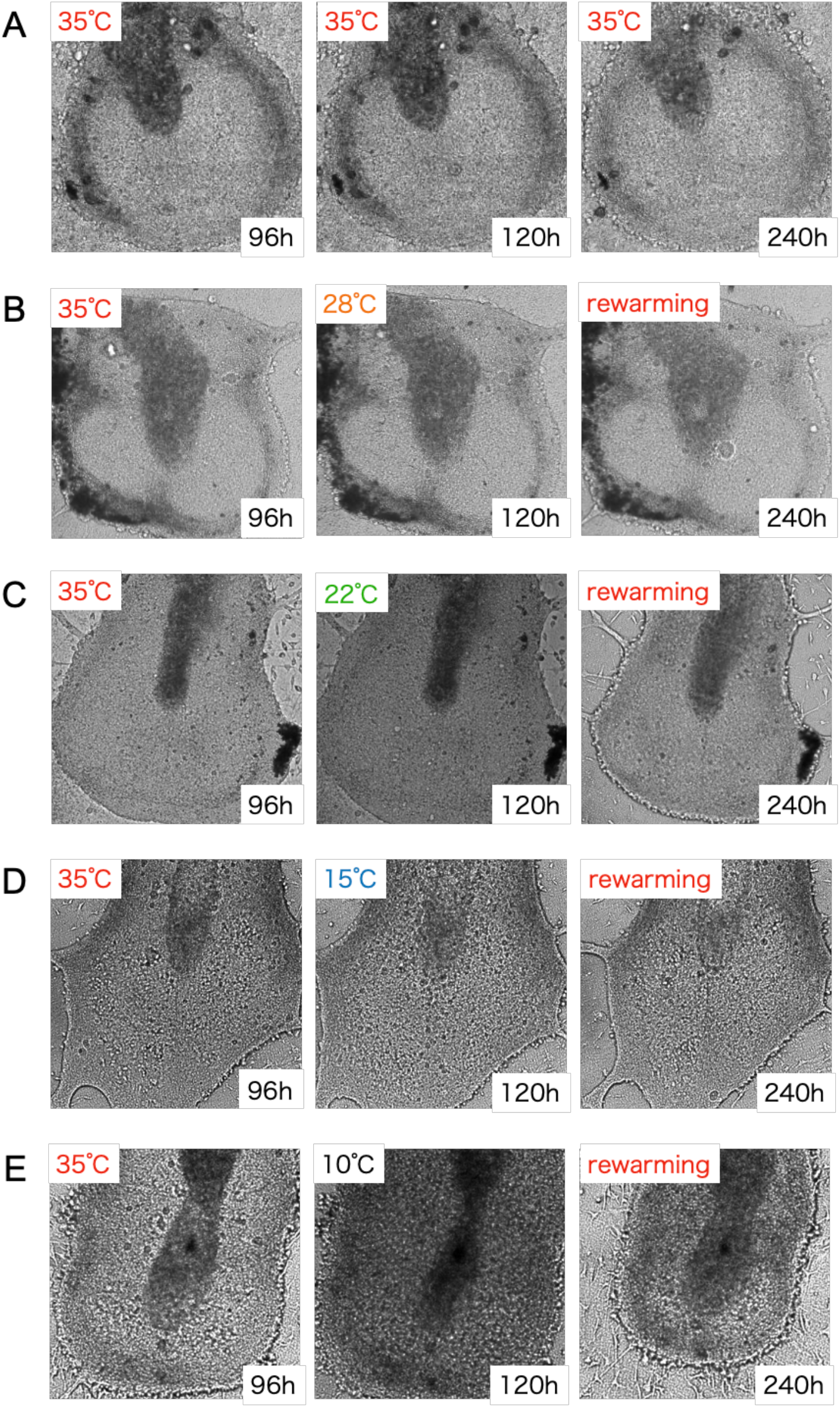
Cold-induced damage in mouse SCN slices. Bright-field images before (left), during (center), and after (right) cold exposure. From top to bottom: 35°C, 28°C, 22°C, 15°C, and 10°C exposure, as indicated in the top left corner. The elapsed time after the start of recording is shown in the lower right corner.

**Supplemental Figure 2.**
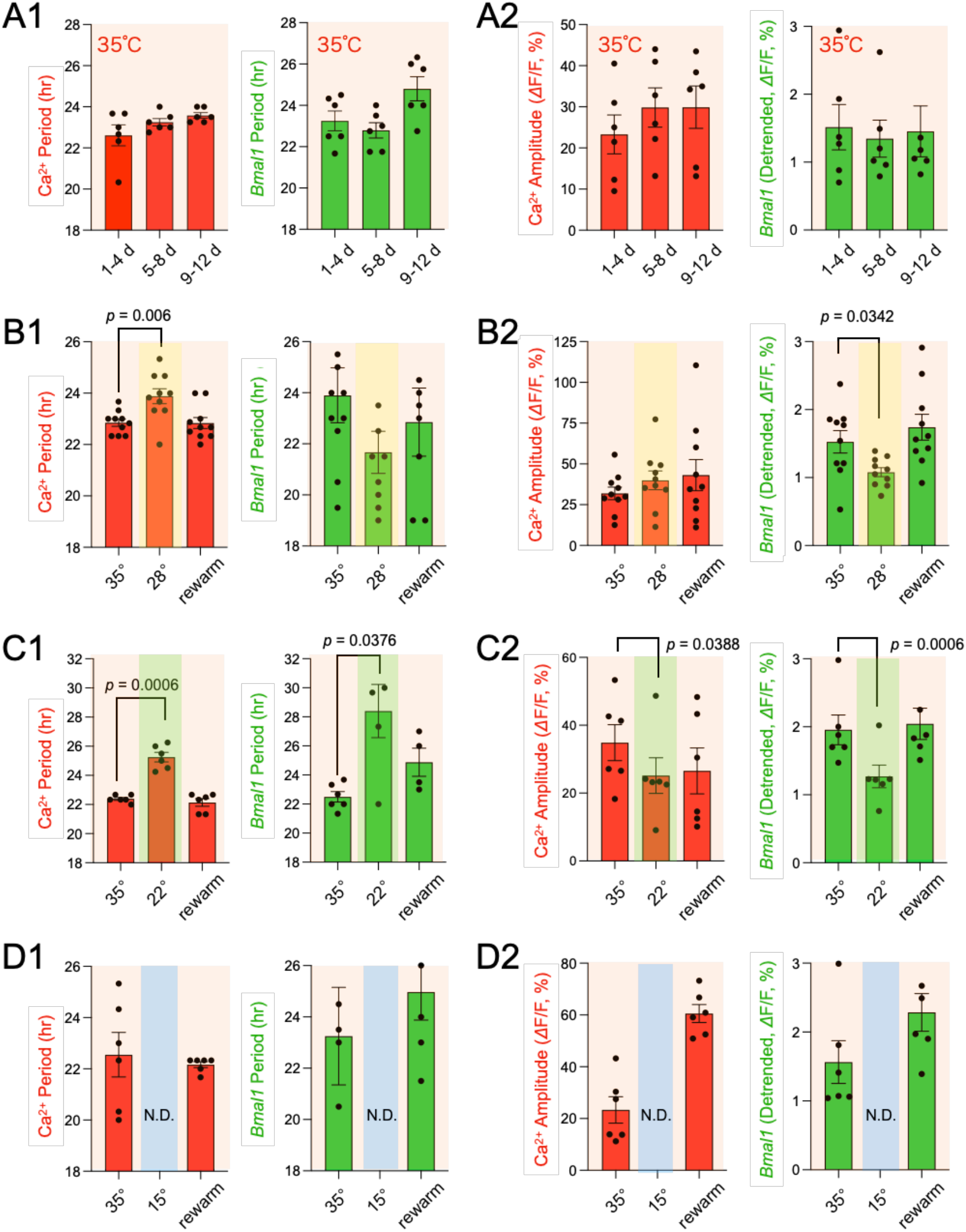
Statistical comparisons of Ca^2+^ and *Bmal1* rhythm periods and amplitudes at various temperatures. Amplitudes (A1, B1, C1, and D1) and periods (A2, B2, C2, and D2) of Ca^2+^ (red, left) and *Bmal1* (green, right) rhythms. Graphs from top to bottom: 35°C (A), 28°C (B), 22°C (C), and 15°C (D). The data are presented as the mean ± SEM.

**Supplemental Figure 3.**
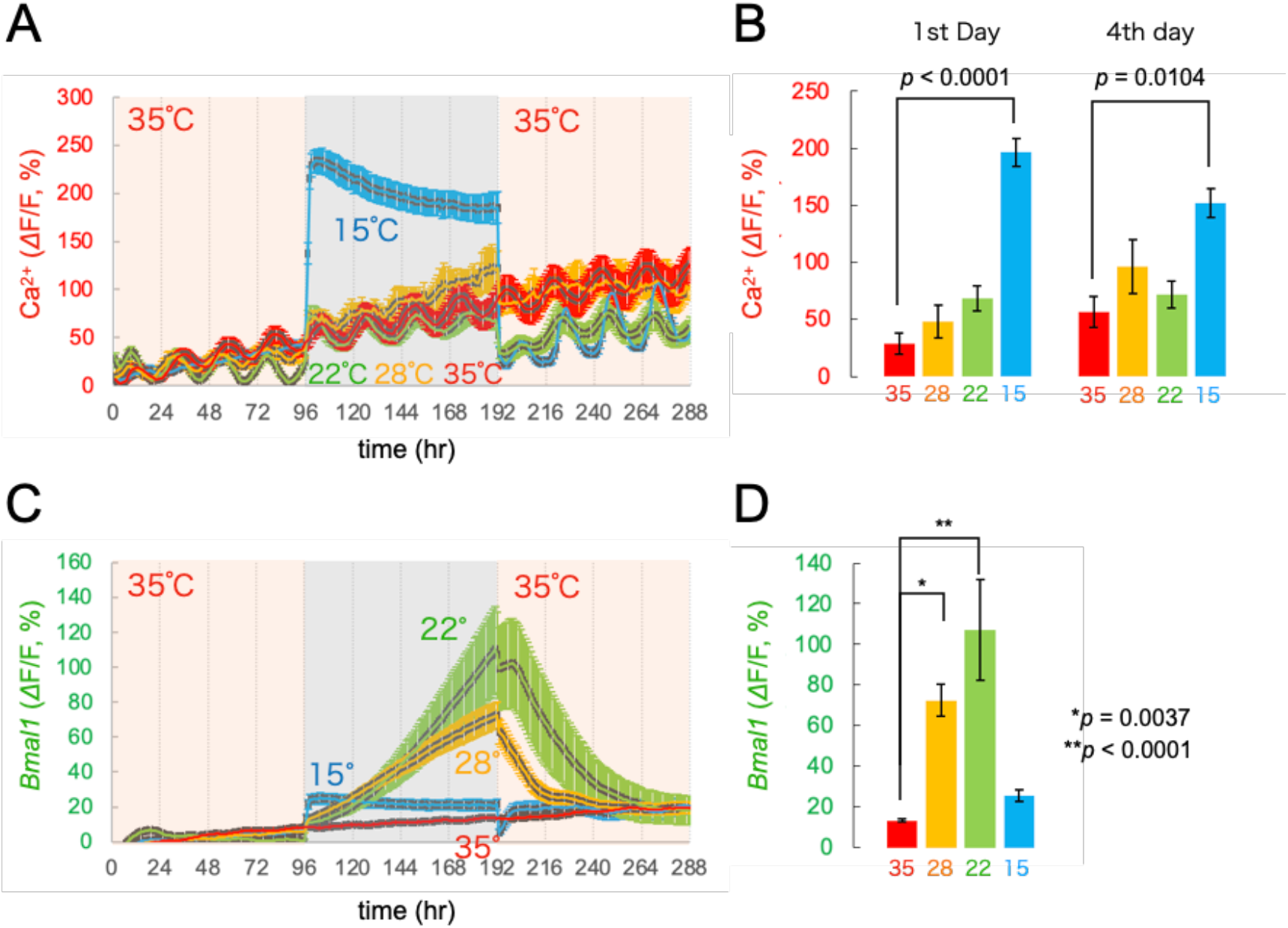
Baseline levels of Ca^2+^ and *Bmal1* signals at various temperatures in SCN neurons. (A) Baseline Ca^2+^ levels at 35°C, 28°C, 22°C, and 15°C. (B) Statistical comparison of baseline Ca^2+^ levels on Days 1 and 4 during cold temperatures. (C) Baseline levels of *Bmal1* signal at 35°C, 28°C, 22°C, and 15°C. (B) Statistical comparison of the *Bmal1* signal maximum baseline levels during cold temperatures. Time is depicted after the start of the recording. All data are presented as the mean ± SEM.

**Supplemental Figure 4.**
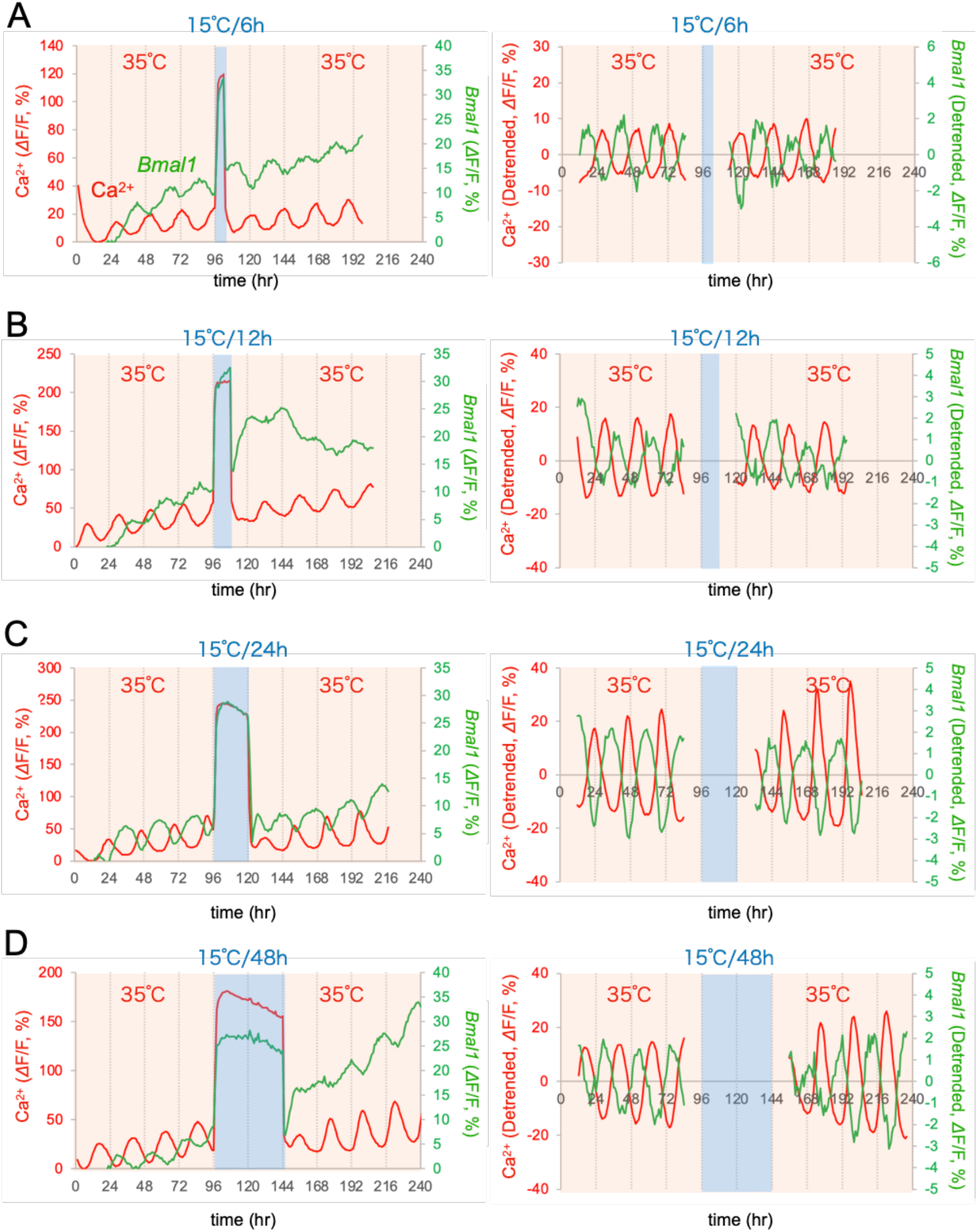
Representative traces of Ca^2+^ and *Bmal1* rhythms in SCN neurons with varying cold exposure times. Representative row (left) and detrended (right) traces of *ΔF/F* (%). After recording Ca^2+^and *Bmal1* rhythms for 4 days at 35°C, SCN slices were exposed in cold for 6 h (A), 12 h (B), 24 h (C), and 48 h (D) at 15°C, followed by rewarming to 35°C for 4 days. All traces are averages in the SCN region. Time is depicted after the start of the recording.

**Supplemental Figure 5.**
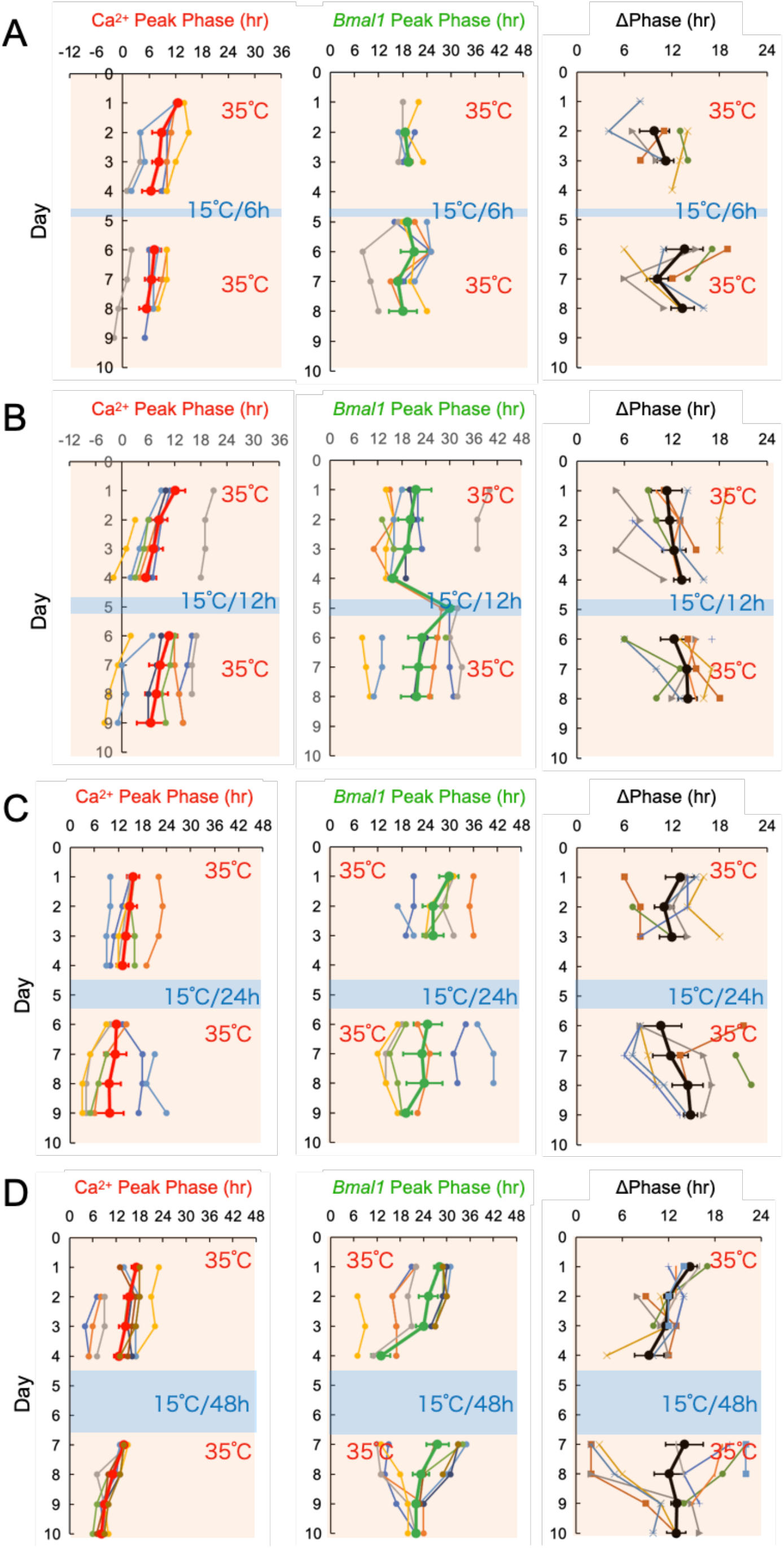
The plots of the peak phases of the Ca^2+^ and *Bmal1* rhythms and phase difference in SCN neurons with varying cold exposure times. After recording Ca^2+^ and *Bmal1* rhythms for 4 days at 35°C, mouse SCN slices were cold-exposed for 6 h (A), 12 h (B), 24 h (C), and 48 h (D) at 15°C, followed by rewarming to 35°C for 4 days. All traces are averages in the SCN region. The data are presented as the mean ± SEM. Time is depicted after the start of the recording.

**Supplemental Figure 6.**
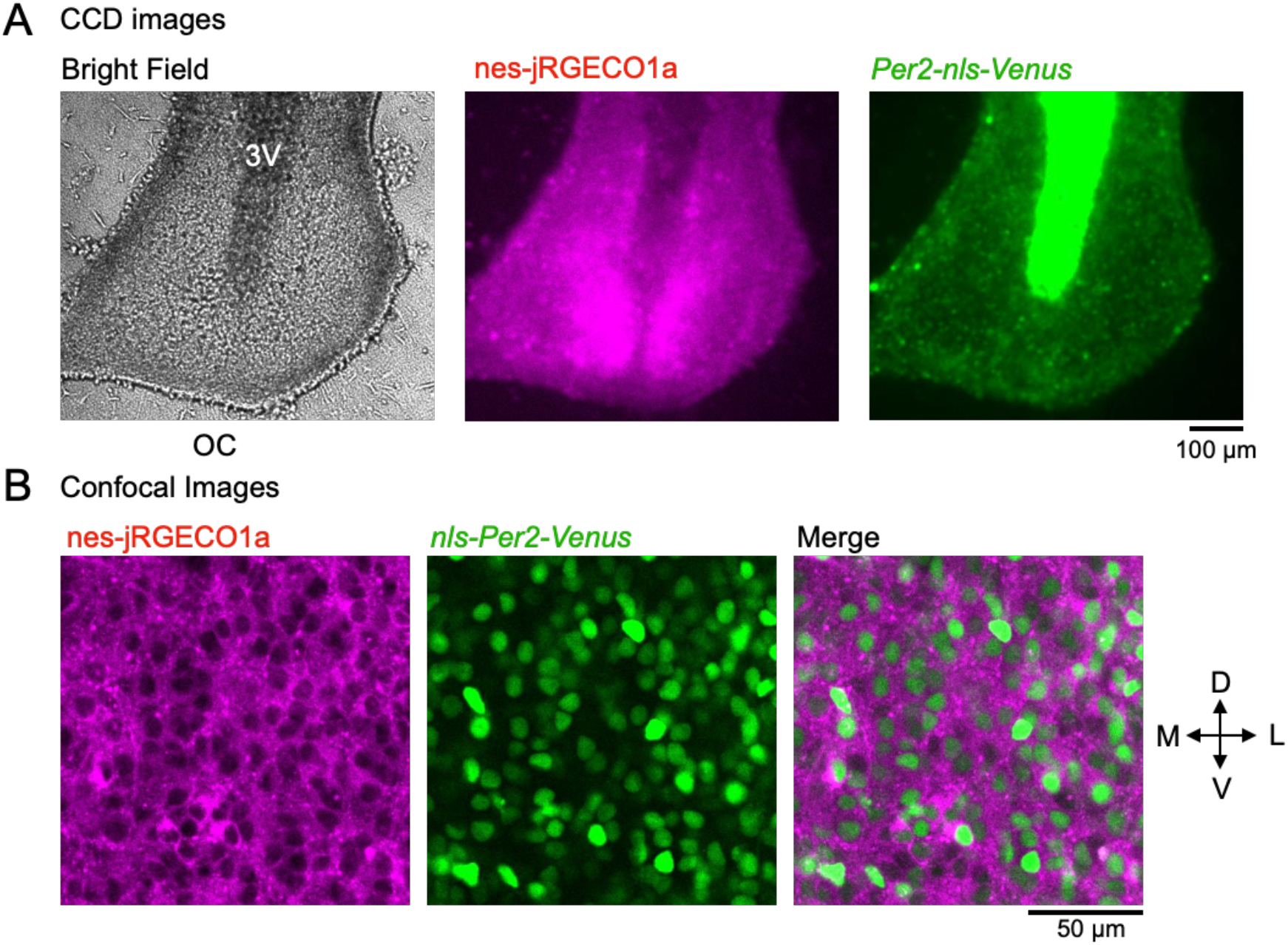
Expression of nes-jRGECO1a and *Per2-nls-Venus* in mouse SCN slices. (A) Representative images of mouse SCN slices expressing nes-jRGECO1a and *Per2-nls-Venus*. Bright-field (left), nes-jRGECO1a (center), and *Per2-nls-Venus* (right). (B) Confocal images of nes-jRGECO1a (left), *Per2-nls-Venus* (center), and merged images on the left side of the SCN slices. The center region of the left SCN is depicted. 3V: the third ventricle, OC: optic chiasma.

